# Molecular phylogenetic analysis for *Ovis aries* with whole mitochondrial genome sequencing

**DOI:** 10.1101/2022.07.29.502105

**Authors:** Aruna Pal, Samiddha Banerjee, Kanai Pathak, Manas Kumar Das, P.N. Chatterjee

## Abstract

Domestication of *Ovis aries* has taken place time immorial across the different geographical regions across the globe. Biodiversity among the sheep population has been interesting to study by a group of researchers based on mitochondrial genes like Cytochrome B, D loop. The current study is a novel attempt to understand the molecular phylogenomics among Ovis aries through all the 37 genes of mitochondria. We have analyzed complete mitochondrial genome sequencing for sheep breeds for West Bengal as Garole, Chotanagpuri, Bonpala and Birbhum sheep. Phylogenetic analysis reveals genetic similarity between Garole and Chotanagpuri brred of sheep, where as Birbhum and Bonpala were found to be genetically distinct. Phylogenomics in a global prospect reveal three lineages, Lineage A comprise of sheep from West Bengal grouped with Tibetan sheep. Lineage B consists of sheep population across the other parts including Europe (France, Denmark), Africa and Syro-Arabian desert, whereas Lineage C emerged as smaller outgroup.

## Introduction

*Ovis aries* is distributed across the globe. It is interesting to understand the molecular evolution of sheep. The sheep are mainly reared for mutton production or wool production. Some of the countries like Australia, sheep industry has a major role in the economy. In developing countries like India, extensive genetic diversity has been noted^**1**^ (kolte paper). Indian sheeps are mostly reared for mutton production and carpet wool production. Apparel wool production is also evident in some sheep breeds. We have studied the biodiversity among the sheep and goat breeds through biomorphometric data analysis and also with mitochondrial genes as Cytochrome B gene^**2**^. Certain other researchers have also studied molecular evolution for sheep based on mitochondrial gene as Cytochrome B and D loop^**1–7**^. Apart from mitochondrial genes, group of Scientists have studied molecular evolution of sheep based on SNPs^**8**^ or through microsatellite marker^**9**^.

In our lab, we had been studied sheep based on its phenotypic and molecular characterization^**2**^ and disease resistance against *Haemonchus contortus*, a dreadly parasite^**10,11**^. We could explore certain uniques genes from these indigenous sheep population that confer molecular resistance against *H. contortus*^**10,11**^. We could equally explored certain mutations in mitochondrial gene as Cytochrome B, which was responsible for sheep disease and debility^**12**^.We had characterized and explored differential mRNA expression profile for indigenous sheep breed based on certain immune response genes as CD4, CD8, Cytochrome B, RIGI, CD14, MD2, LBP, NFkB, IL12, MyD88, IL6, IL1B, IL10, TNFalpha, TLR4^**10–15**^. We had explored and documented a new sheep breed from Birbhum district of West Bengal, and declared as “Birbhum sheep” ^**16**^. We had studied mitochondrial genes from other livestock as pig^**17,18**^ and poultry as duck^**19–21**^ and goose^**21–22**^.

Whole mitochondrial genome map reveals 37 genes with thirteen polypeptide genes, RNA genes and control region as D loop. It is better to study molecular evolution with mitochondrial genes, since they lack intron and are maternally inherited. In our lab, we have studied whole mitochondrial genome sequencing for Ghungroo pig^**18**^, Bengal duck^**19–21**^ and Goose^**21–22**^.

In this current study, we aim to study the molecular evolution of indigenous sheep of West Bengal namely Garole, Birbhum, Bonpala and Chotanagpuri with the help of Whole mitochondrial genome sequencing, and its comparison with WMGS of sheep population globally.

## Materials and Methods

### Whole mitochondrial genome sequencing

Whole mitochondrial genome analysis (WMGA) was conducted for Garole, Birbhum, Bonpala and Chotanagpuri sheep from West Bengal of India. The mitochondrial map was generated. The steps for next generation sequencing studies (Whole mitochondrial genome sequencing) involves isolation of mtDNA, qualitative and quantitative analysis of g-DNA: PCR amplified with COX-2(mt specific), GAPDH and Beta actin primers to validate. The next step involves preparation of library: The paired-end sequencing library (NEBNext Ultra DNA Library Preparation Kit), quantity and quality check (QC) of library on Bioanalyzer: Bioanalyzer 2100 (Agilent Technologies) using High Sensitivity (HS) DNA chip. The final step is cluster Generation and Sequencing. The adapters are designed so as to allow selective cleavage. Whole mitochondrial genome sequences for Ovis aries were retrieved from other parts of the world.

### Construction of Phylogenetic tree

Phylogenetic tree was constructed based on neighbour joining method with Jukes-Cantor substitution model of MAAFT software^**23**^ with all the available sequences from gene bank. We generated phylogenetic tree among the sheep population of these countries.

### Comparison of mitochondrial gene sequences for Indian sheep with sheep from other parts of the world

Nucleotide sequences such otained was subjected to multiple sequence alignment MAFFTsequences. Alignment report was visualized in MSA viewer. SNPs were identified through Lasergene DNASTAR software^**24**^.

### Comparative analysis for amino acid sequences for the coding genes for Indian sheep with respect to others

Derieved amino acid sequences for all the 13 coding genes for mitochondrial genes were aligned for the detection of amino acid sequence variabilities.

### Three-dimensional structure prediction and Model quality assessment

Accordingly in the next step, 3D structural analysis for the polypeptide (13 in numbers) were predicted. The templates which possessed the highest sequence id entity with our target template were identified by using PSI-BLAST (http://blast.ncbi.nlm.nih.gov/Blast). The homology modeling was used to build a 3D structure based on homologous template structures using PHYRE2 server^**25**^. The 3D structures were visualized by PyMOL (http://www.pymol.org/) which is an open-source molecular visualization tool. Subsequently, the mutant model was generated using PyMoL tool. The Swiss PDB Viewer was employed for controlling energy minimization. The structural evaluation along with a stereochemical quality assessment of predicted model was carried out by using the SAVES (Structural Analysis and Verification Server), which is an integrated server (http://nihserver.mbi.ucla.edu/SAVES/). The ProSA (Protein Structure Analysis) webserver (https://prosa.services.came.sbg.ac.at/prosa) was used for refinement and validation of protein structure^**26**^. The ProSA was used for checking model structural quality with potential errors and the program shows a plot of its residue energies and Z-scores which determine the overall quality of the model. The solvent accessibility surface area of the genes was generated by using NetSurfP server (http://www.cbs.dtu.dk/services/NetSurfP/)^**27**^. It calculates relative surface accessibility, Z-fit score, the probability for Alpha-Helix, probability for beta-strand and coil score, etc. TM align software was used for the alignment of 3 D structure of IR protein for different species and RMSD estimation to assess the structural differentiation^**28**^. The I-mutant analysis was conducted for mutations detected to assess the thermodynamic stability. Provean analysis was conducted to assess the deleterious nature of the mutant amino acid. PDB structure for 3D structural prediction of gene for duck was carried out through PHYRE software^**25**^. Protein-protein interaction have been studied through String analysis^**29**^, along with the details of the bioinformatics software^**30,31**^.

## Result

### Characterization of whole mitochondrial genome for Sheep breeds

Mitochondrial map for complete mitochondrial sequence for sheep breeds as Garole Ac no. (Fig 1),Birbhum (Fig 2), Bonpala (Fig3) and Chotanagpuri (Fig4). Whole mitochondrial genome sequences for sheeps from other parts of the world were retrieved from gene bank.

**Figure 1.**
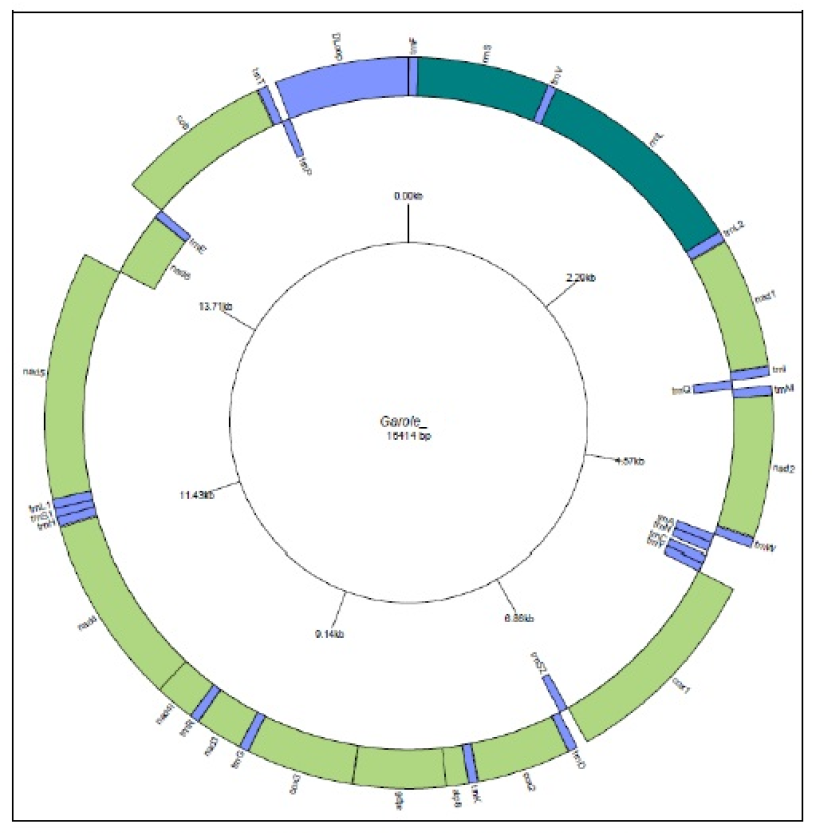
Circular map of Garole mitochondrial genome drawn using GenomeVx representing 37 protein coding genes with total genome size of 16,414 bp.

**Figure 2:**
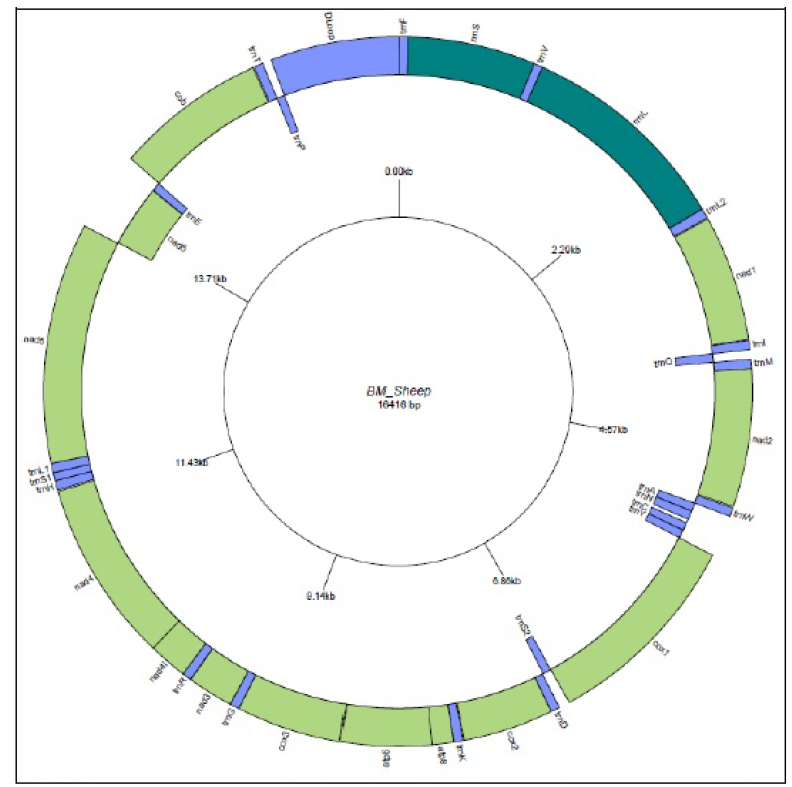
Circular map of BM_Sheep mitochondrial genome drawn using GenomeVx representing 37 protein coding genes with total genome size of 16,416 bp.

**Figure 3.**
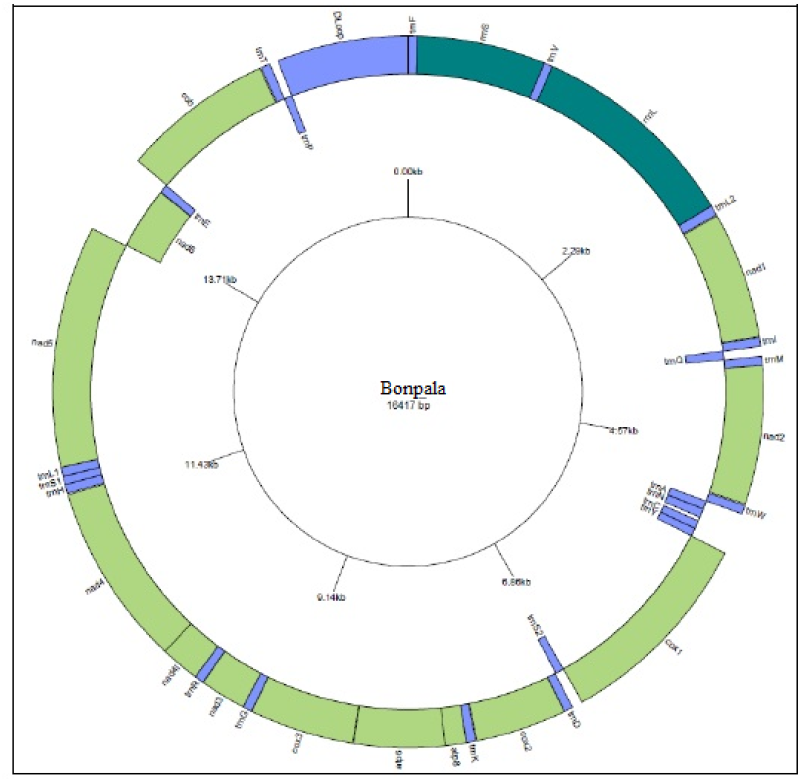
Circular map of Bonpala mitochondrial genome drawn using GenomeVx representing 37 protein coding genes with total genome size of 16,417 bp.

**Figure 4:**
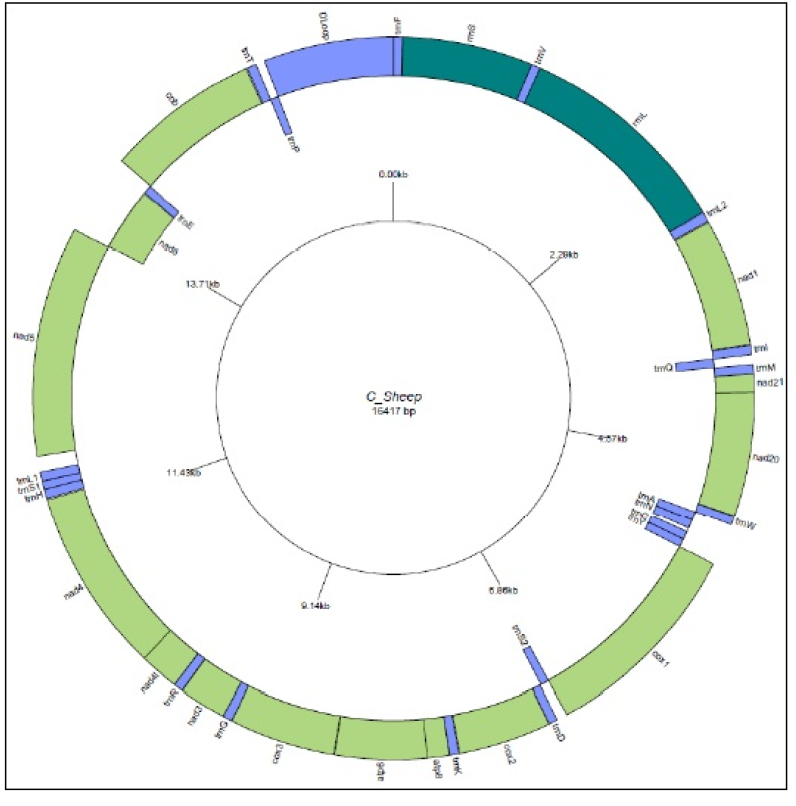
Circular map of C_Sheep mitochondrial genome drawn using GenomeVx representing 37 protein coding genes with total genome size of 16,417 bp.

### Construction of Phylogenetic tree

Phylogenetic tree was constructed based on neighbour joining method with Jukes-Cantor substitution model of MAAFT software with sheep of West Bengal as represented in Fig 5.

**Fig 5:**
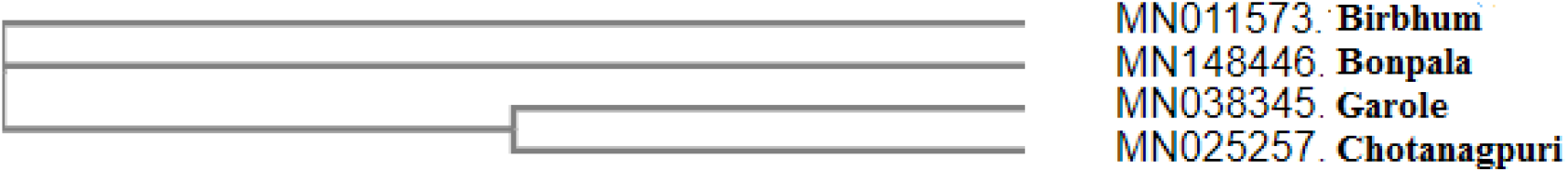
Phylogenetic tree for Sheep breeds from West Bengal, India

### 3D structural analysis for derived peptides for the coding genes of mitochondria

The polypeptide sequence from derived complete mitochondrial genome sequence for sheep breed as Garole, Bonpala and Birbhum were compared (Fig 6). Certain SNPs were identified among the nucleotide sequences. But the most interesting fact to note is that the mutations were non-synonymous without amino acid differences detected.

**Fig 6:**
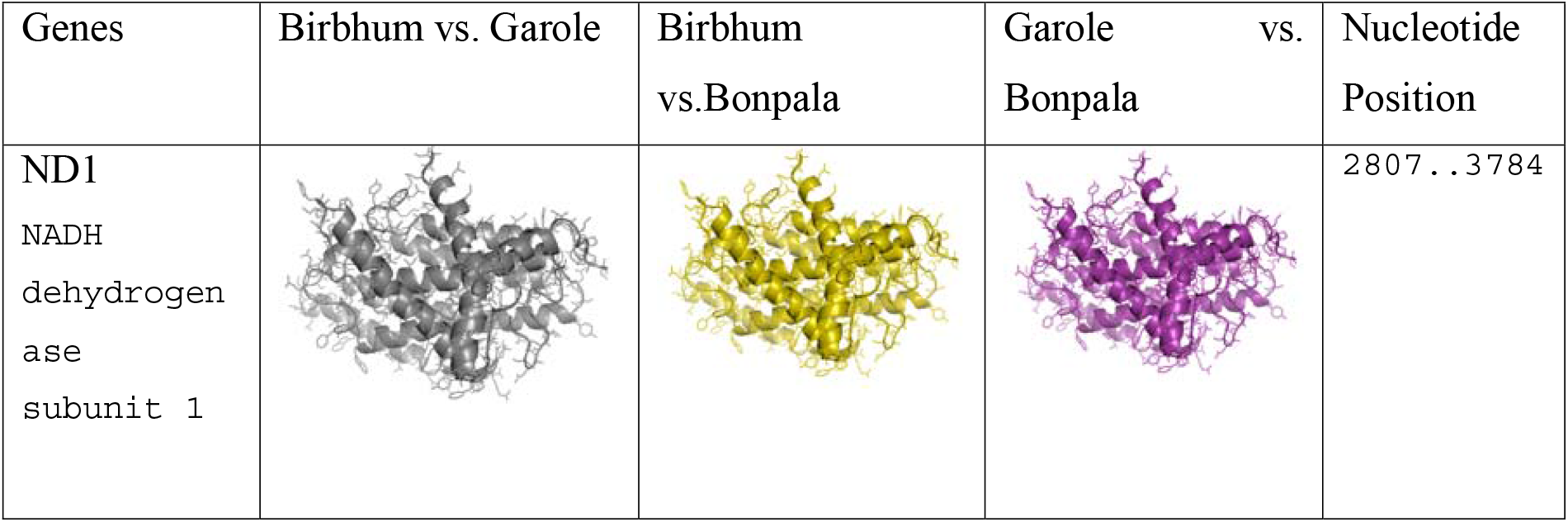
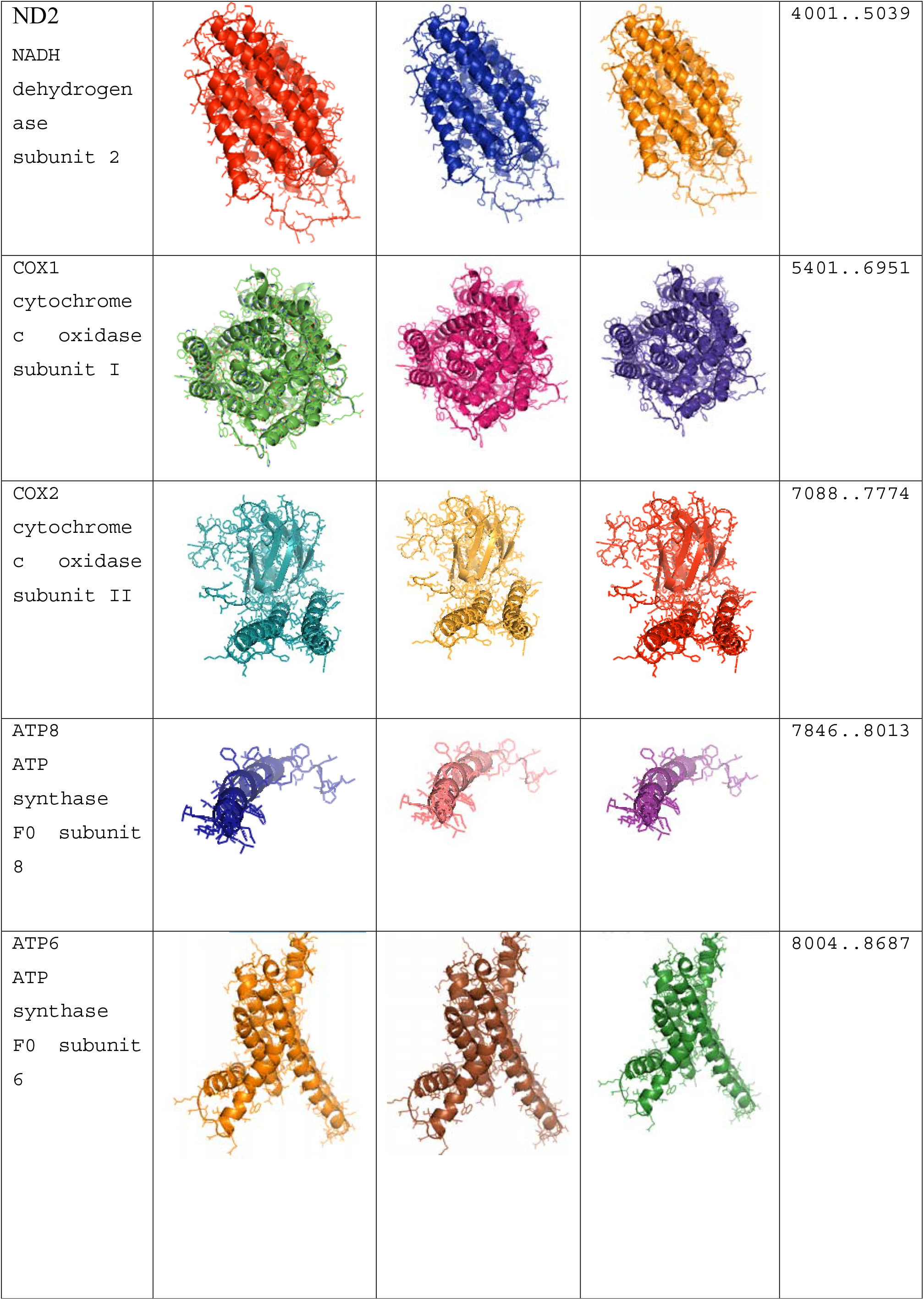
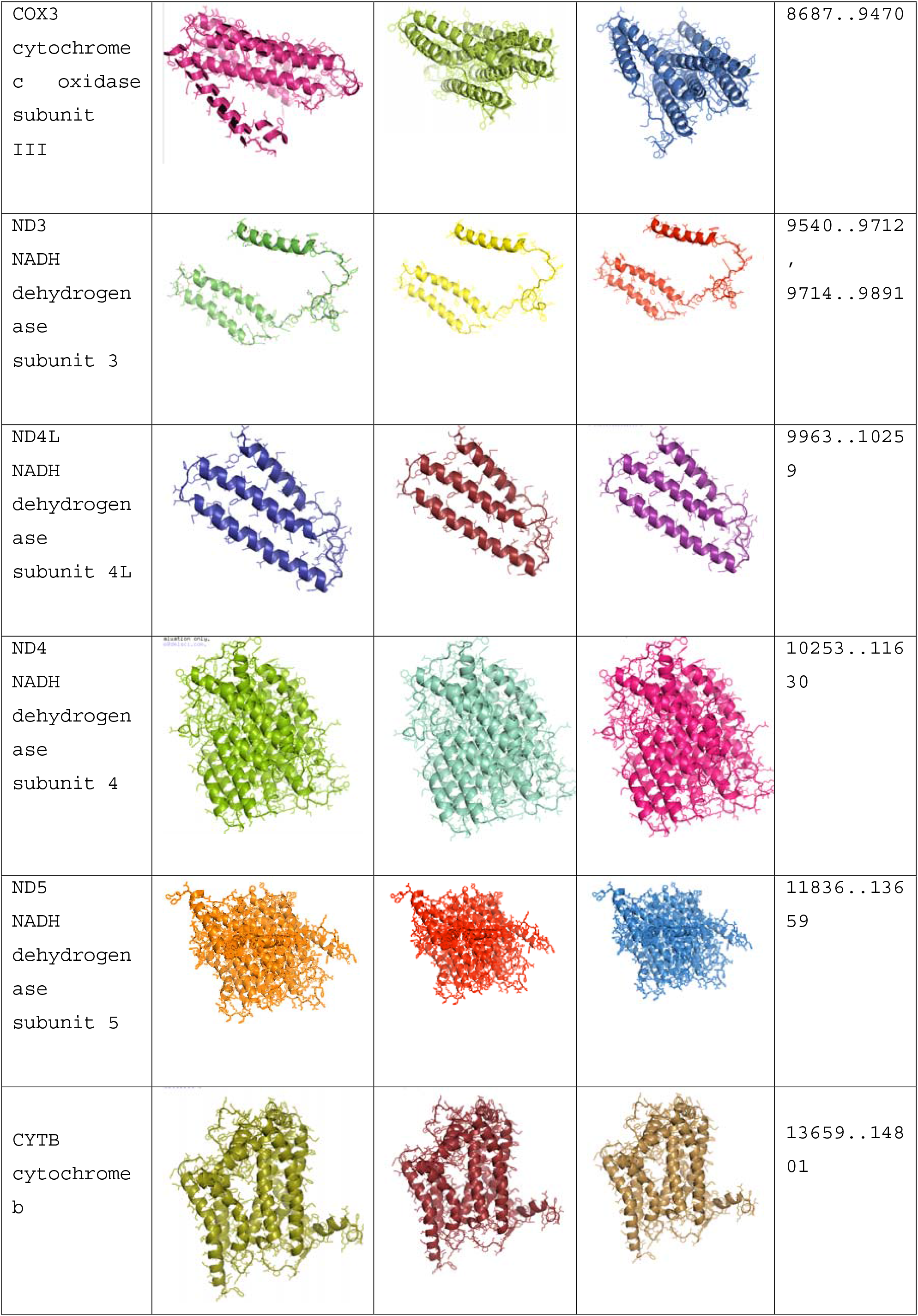
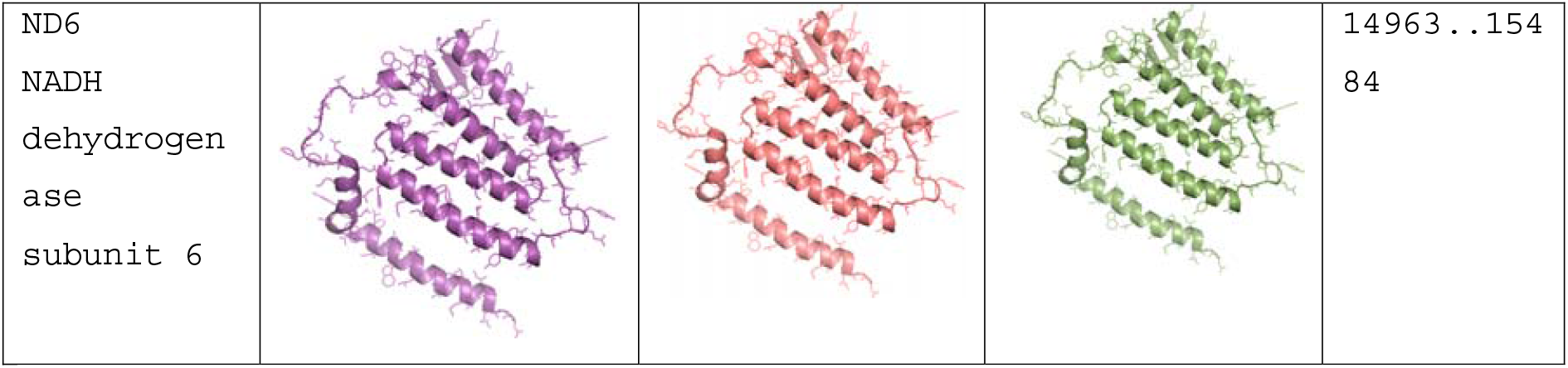
Comparison of derived peptides for mitochondrial genes for Birbhum sheep w.r.t Garole and Bonpala

The whole mitochondrial genome sequencing reveals 37 genes with 13 coding genes, non coding genes and control D-loop region. 3D structural analysis of derieved amino acids with respect to 13 polypeptide genes were conducted. Although sequence variabilities were observed at nucleotide level, aminoacid sequenceforthese 13 coding genes for mitochondria were observed to be conserved across the globe. Alignment for the 3D structure for these coding genes were found to have RMSD as zero, implying no structural variation (Fig6). The molecular interaction for mitochondrial gene has been depicted in figure 7.

**Fig 7:**
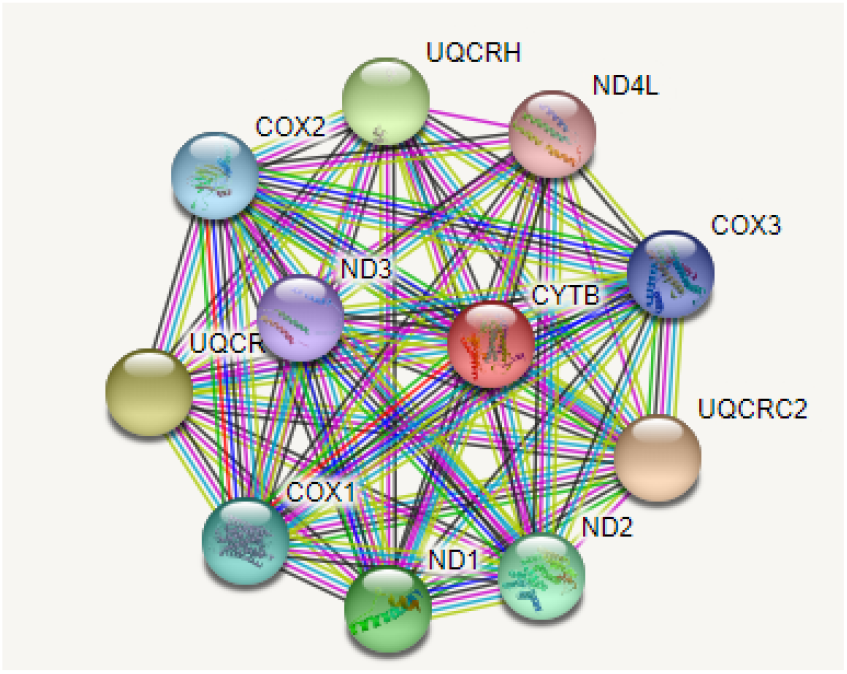
Molecular interaction among the mitochondrial gene in *Ovis aries*.

### Molecular phylogenetic analysis for sheep population reared globally

In the current study, we have analyzed phylogenetic relationship of sheep globally (Fig 8). We have sequenced the complete mitochondrial genome for sheep breeds from West Bengal, India. We had retrieved complete mitochondrial genome sequences for sheep breeds reared across the world from gene bank. We observed distinct breed identity among the sheep breeds. Birbhum sheep was observed to be genetically closer to Tibetan sheep(KU575248). On the otherhand, we observed genetic closeness of Bonpala sheep with Sishui fur (MW364895) and other breed (MT768239.1) from China. Three distinct lineages were observed. The lineage A contain contains Indian sheep along with other Asian countries as China.Lineage B contains European breed as Rambouillet sheep, sheep from France, Africa and Arab countries. Lineage C is Chinese in origin.

**Fig 8:**
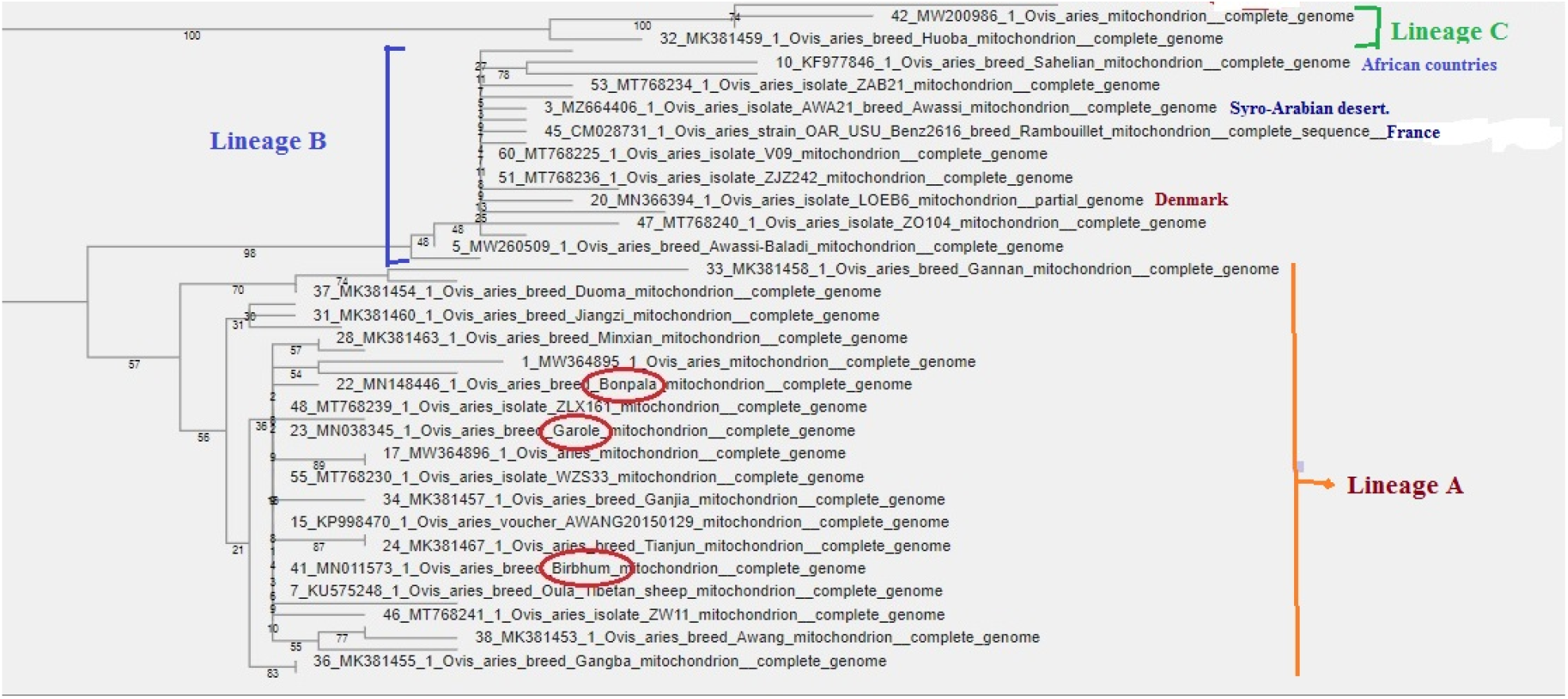
Molecular phylogeny for Ovis aries distributed globally based on Complete mitochondrial genome.

## Discussion

The evolution and domestication of sheep is interesting to study. Domesticated sheep or Ovis aries are reared worldwide. But the purpose of rearing differs from place to place. In some places, they are reared for mutton production, whereas in other parts, they are reared for apparel wool or carpet wool. The geographical distribution for sheep habitat ranges from temperate region, tropical region, dry arid region and also coastal region, where they are highly adapted and resistant to common diseases. As a result varied biodiversity among sheep population is evident.It has been reported that sheep and dog were the first livestock to be domesticated^**32**^.

Evolution of sheep has been studied by a number of scientists across the globe based on D-loop, ND or Cytochrome B gene as mitochondrial gene^**1–9**^. The current study revealed that mitochondrial genes with all 37 genes including 13 coding genes and rest non coding and a control region (D loop), provide a better in site for evolutionary studies. The current study is unique in the sense that it employs all 37 mitochondrial genes for studying evolutionary studies for *Ovis aries* across the globe, instead of few genes for the first time.

In the current study, we had analyzed whole mitochondrial genome sequencing for the indigenous sheep. The mitochondrial genome consists of 13 protein-coding genes, 2 rRNA genes (12S rRNA and 16S rRNA), 22 tRNA genes, and 1 control region (CR). The total length of the genome is 16,623⍰bp, with a base composition of 29.66% A, 22.28% T, 15.35% G, and 32.71% C. The total length of 13 protein-coding genes is 10,997⍰bp long, all of which are encoded on the same strand except for ND6 in the light strand. Except for ND2, ND5 and Cytb (ATC start codon), ND6 (TAA start codon), and COX1,COX2 (GTG start codon), the remaining 7 protein-coding genes initiate with ATG (COX3, ND1, ND3, ND4, ND4L, ATP6, ATP8). and retrieved the gene bank sequences for complete mitochondrial genome from *Ovies aries* across the globe. Mitochondrial genome map comprises of 37 genes with 13 polypeptide coding genes. We had also observed from string analysis that there is interaction among these mitochondrial genes. As a result the comprehensive study involving all mitochondrial genes provides us with better research output.

We had revealed distinct breed identity for sheep population from West Bengal, India as Garole, Birbhum, Bonpala and Chotanagpuri based on Whole mitochondrial genome sequencing. Garole breed of sheep was observed to be genetically closer to Chotanagpuri breed. Earlier, in our lab, we had reported similar result with Cytochrome B sequence analysis^**2**^ for these sheep breeds.Similar distinctness for these breeds have been reported through microsatellite analysis^**16**^. Studies on sheep evolution and hylogenetic analysis have been studied across the globe based on either single or two mitochondrial gene, microsatellite, SNP or Y chromosome, but phylogenetic analysis through complete mitochondrial genome sequences are reported here for the first time. Molecular phylogeny for sheep have been studied in different countries as Nepal^**33**^, India^**34,1**^, China^**9**^, Tibetan sheep^**5**^, Europe^**4,8**^, Kenya^**6**^, Bangladesh^**7**^.

In this current study we had identified distinct lineages of sheep population, belonging to different clades and haplotype distributed world wide in mostof the above discussed studies^**1,4,8,34**^. One of the most interesting thing observed was that Indian sheep population were grouped in Lineage A. Chinese sheep population, particularly Tibetan sheep were the closest ancestor to Indian sheep breeds, particularly that of West Bengal.

## Acknowledgement

The authors would like to thank Department of Biotechnology, Ministry of Science and Technology, Govt. of India for providing the financial support for carrying out the research work (Grant No. BT/Bio-CARe/04/10100/2013-14). The funders had no role in study design, data collection and analysis, decision to publish, or preparation of the anuscript. The authors are equally thankful to West Bengal University of Animal and Fishery Sciences for carrying out the work.

